# Mining functional annotations across species

**DOI:** 10.1101/369785

**Authors:** Sven Warris, Steven Dijkxhoorn, Teije van Sloten, Bart van de Vossenberg

## Abstract

**Motivation:** Numerous tools and databases exist to annotate and interpret the functions encoded in genomes (InterProScan, KEGG, GO etc.). However, analyzing and comparing functionality across a number of genomes, for example of related species, is not trivial.

**Results:** We present a novel approach, for which KEGG and Gene Ontology data are imported into a Neo4j graph database and InterProScan results from several species are added. Using the Neo4j plugin for Cytoscape, users can query this database and visualize functional annotations (sub)graphs, to compare and group functional annotation across species.

## Introduction

A first step following *in silico* reconstruction of a genome[1] is usually to produce a structural annotation of the genome to identify repeats[2], genes[3,4] and other genomic features[5]. Combined with sequence information, these data can then be used for comparative genomics efforts to interpret the functional makeup of the organism under study to that of a related organism. Genes can be functionally annotated using InterProScan[6] or Blast2GO[7], studying presence/absence of specific Kyoto Encyclopedia of Genes and Genomes (KEGG)[8] pathway elements[9] or Gene Ontology (GO) enrichment[10]. These approaches mainly focus on comparing a single species of interest to one known reference species. However, there are currently no tools available to store, visualize and analyze the functional annotation of several species combined. Here, we present a new approach building on graph database technologies[11] and Cytoscape[12] to mine functional annotations, enable comparison of GO terms and KEGG pathways and visualize the results[13].

Both GO and KEGG databases store highly connected information, suitable for storing in a Neo4j graph database [14]. After addition of InterProScan results, these graphs can be relatively easily queried to perform complex analyses on the data. The applicability of this approach is demonstrated, by analyzing seventeen fungal species, including two *Synchytrium endobioticum* strains and six reference species, divided into four functional groups [Vossenberg & Warris et al., unpublished]. Here we outline the methodology underlying the approach.

## Materials and methods

The data set consists of the genes of seventeen fungal species, including six reference species, in total 157,709 genes. The seventeen species were divided into four groups: ‘Chytrids obligatoire biotroof’ (ChytObl), ‘Chytrids culturable’ (ChytCult), ‘Control obligatoire biotroof’ (CtrlObl) and ‘Control culturable’ (CtrlCult). InterProScan version 5.16‐55.0[6] was used to functionally annotate the genes.

### Software and database

Python version 3[15] and the BioPython package[16] were used for processing the protein files, InterProScan output, the Gene Ontology (GO) database[10] and for connecting to the KEGG API[17]. In particular, the Python modules Bio.KEGG and Bio.graphics were used to connect to KEGG and process the results. The Python package Neo4j V1[18] was used to connect to the Neo4j database using the Bolt network protocol.

## Implementation

### Processing the Gene Ontology data

The GO-basic data set was downloaded from the Gene Ontology Consortium website (http://geneontology.org/), processed with a Python script and stored in a Neo4j graph database, with each node corresponding to a GO term and “ISA” relations as edges. It is important to note that the GO database does not follow a tree structure but rather a more general Directed Acyclic Graph (DAG), i.e. many GO terms have more than one parent. Based on the InterProScan output, the number of times a GO term is found in a certain species is added as a field to the node representing that particular GO term. Hence, after processing all data, each GO term node holds the total number of times it has been assigned to a particular organism. Counts per species group are determined by summing individual species counts. By propagating the counts from the leaves to the three toplevel GO terms “biological process”, “molecular function” and “cellular component”, each level in the GO database contains the number of times that GO term and all its descendants are found by InterProScan. However, as the GO database is a DAG, a descendant can be linked more than once to a level yet count values should only be used once. A query in Cypher, the builtin Neo4j query language (https://neo4j.com/developer/cypherquerylanguage/), selecting only all distinct GO terms below a given level helps prevent double counting (Query 1). Figure 1 shows an example of the toplevel node ‘biological process’ and some of it descendants.

**Figure 1:**
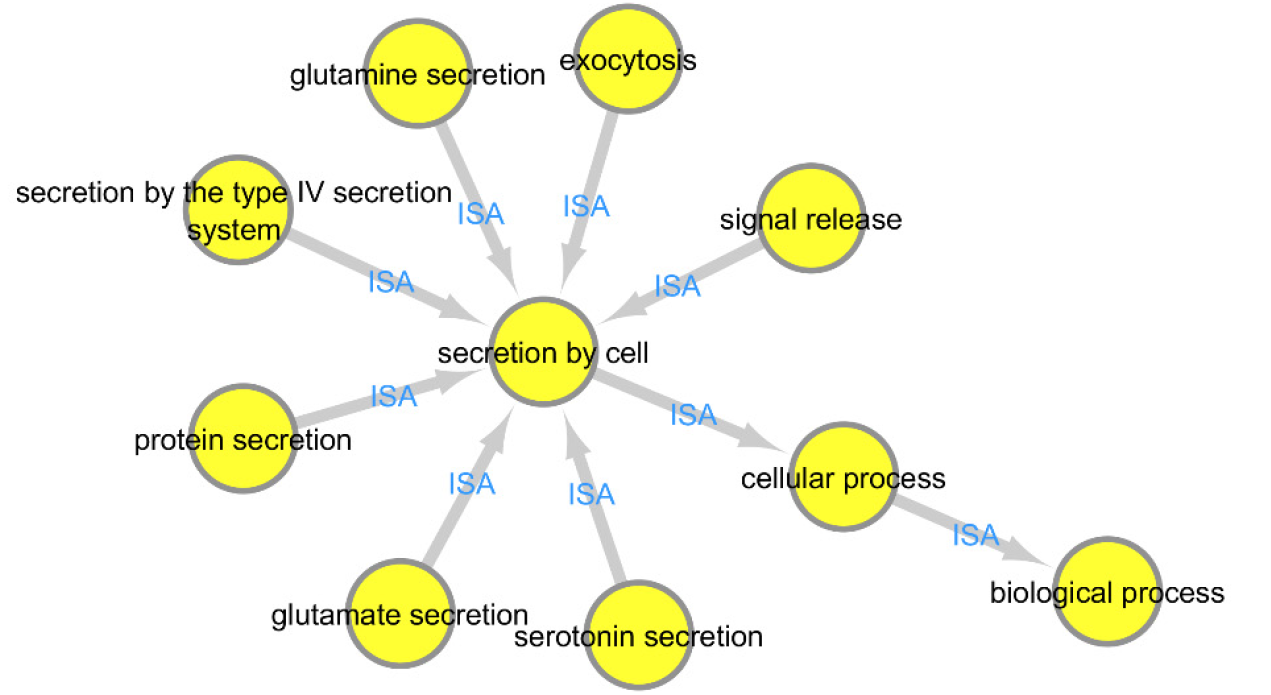
Part of the GO term DAG, showing the toplevel node ‘biological process’ and some of it descendants.

~~~
match (a:GOTerm) with collect (a) as allGo
unwind allGo as goTerm
match (goTerm)<-[r:ISA*]-(b:GOTerm) with collect (distinct b) as allB, goTerm
set goTerm.all{orgName} =
    goTerm.{orgName} + reduce (allUp = 0, n IN allB | allUp + n.{orgName})
~~~

Query 1: Propagating the number of times a GO term was found in the InterProScan results up through the graph. The *collect (distinct b)* part of the query ensures nodes are used counted once.

### KEGG pathways

KEGG pathway identifiers and EC numbers are extracted from the InterProScan output. Using the KEGG API, the structures of the found pathways are downloaded in XML (KGML) format and stored in the Neo4j database. Pathway elements are directly translated into nodes (‘enzyme’, ‘pathway’, ‘map’, etc.) and relationships (‘ECrel’, ‘maplink’, ‘reaction’ etc.). Enzymes are linked to a pathway using the ‘in’ relationship. An enzyme can occur in more than one pathway and can therefore have more than one outgoing ‘in’ relationship. The XML data also contains graphical data to place enzymes on the KEGG pathway images. These graphical properties are set as attributes of the ‘in’ relationship to arrive at a unique enzyme-pathway link. Relations between an enzyme and its substrates and products are not available in the XML data. Therefore, for each of the enzymes the KEGG annotation is retrieved through the API and the relationships ‘usedby’ and ‘produces’ are created to connect enzymes with compounds. The resulting graph is a Directed Cyclic Graph (DCG, Fig. 2).

**Figure 2:**
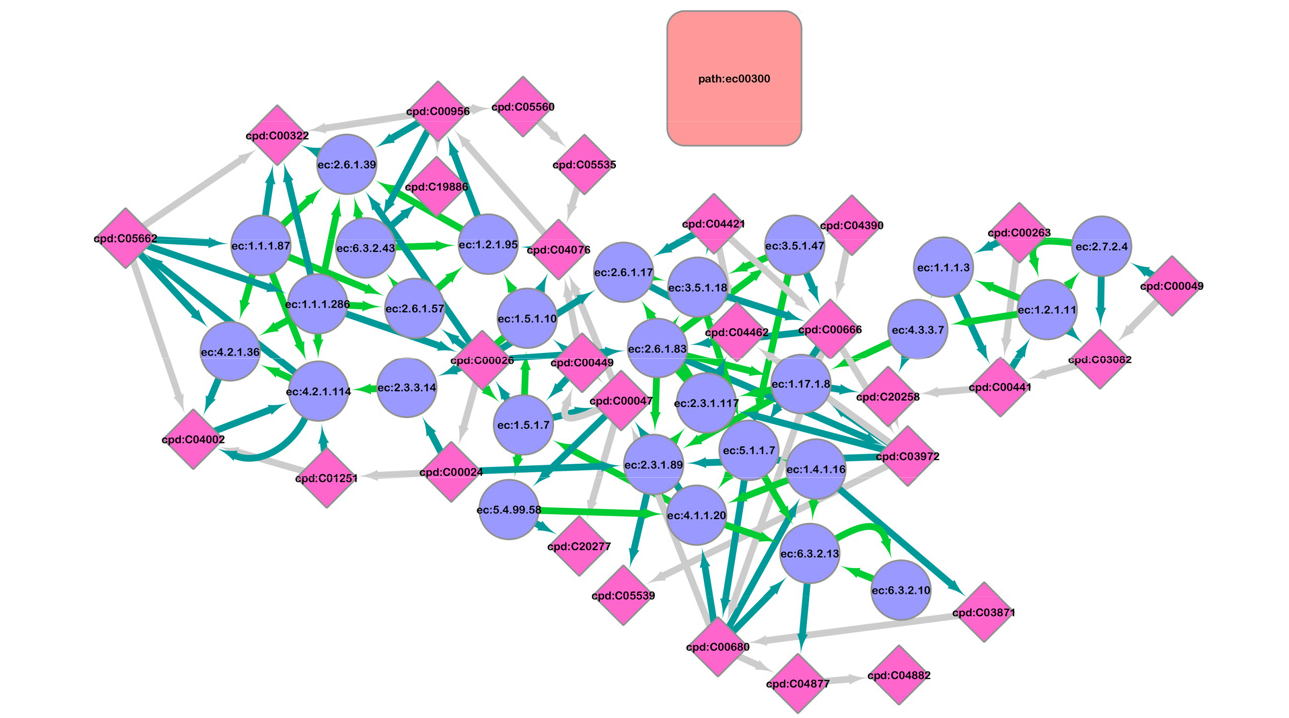
KEGG pathway ec:00300 (pink rounded box) with its enzymes (blue circles) and compounds (pink diamonds). For clarity of presentation, the ‘in’ relationship is not shown.

For each of the seventeen organisms, the number of times an enzyme is predicted by InterProScan is added to the particular enzyme node. Similar as for the GO terms, counts are aggregated for species groups and added to the enzyme nodes. As the KEGG pathway graph is a DCG, no propagation to a higher level can be performed.

### Connecting KEGG to GO

The KEGG database contains links to GO terms for many of its enzymes. Using the same data as for connecting compounds to enzymes, the GO terms are linked to the enzymes with the ‘crossConnect’ relationship (Fig. 3).

**Figure 3:**
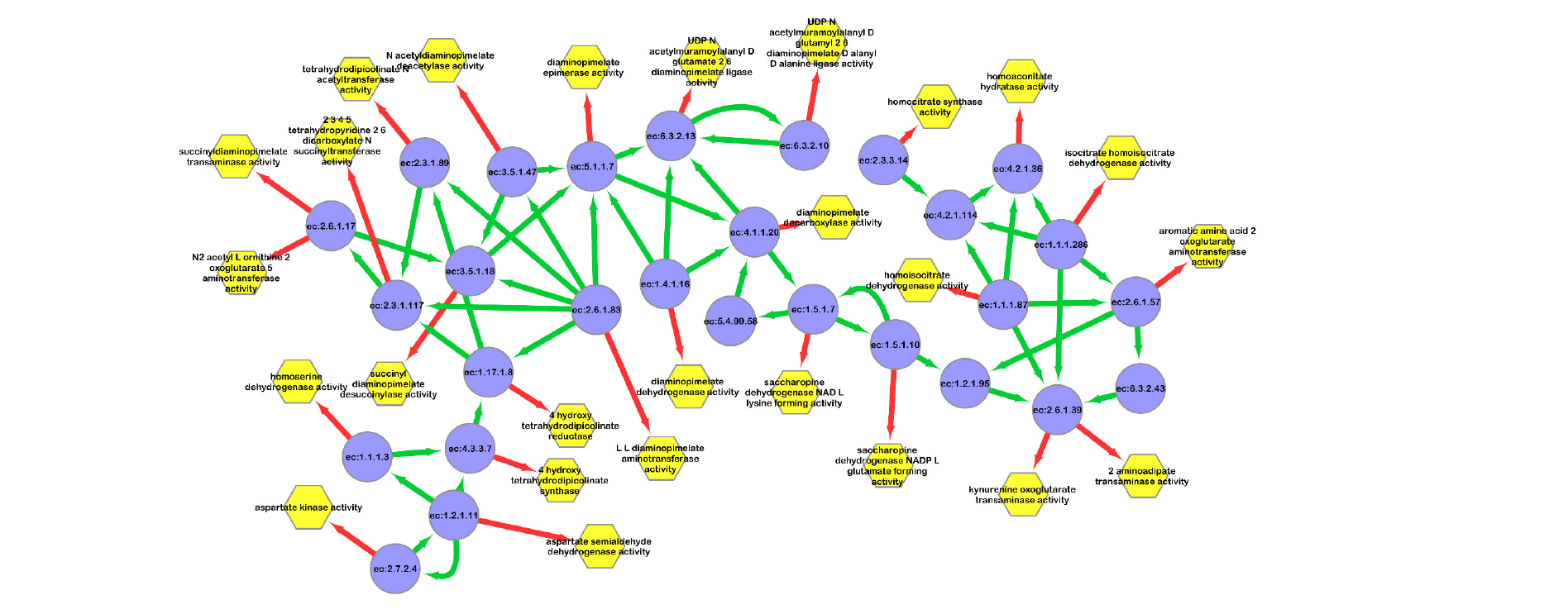
KEGG enzymes (blue circles) and GO terms (yellow hexagons) linked through the ‘crossConnect’ relationship (red edges).

### Cytoscape and the Neo4j plugin

Cytoscape[12] is a commonly used tool for the visualization and analysis of biological networks, such as protein-protein interaction networks[19] and gene coexpression networks[20]. Cytoscape enables the user to define presentation styles based on node and edge attributes, merge networks and visualize the network using a large collection of different layouts. However, loading all data stored in our Neo4j database (Tables 1 and 2) directly is technically impossible due to memory constraints. Moreover, it is visually unattractive: it will produce a complex ‘hairball’ in which nothing is distinguishable. Therefore, an already available Java implementation of a basic Neo4j Cytoscape plugin, cyNeo4j[21], was refactored and extended to create a new plugin to connect Cytoscape directly with a Neo4j database[14]. The plugin allows the user to query the database and store created/imported networks in the database. To allow generic reuse, the plugin also has the ability to read an XML file with predefined Cypher queries. The network resulting from running such a query then becomes available in Cytoscape for visualization; all figures in this manuscript were created using Cytoscape and the Neo4j plugin. The generic forms of these queries are available in XML on the Github page[22]. Additional features of the plugin include exporting networks to Neo4j and interactively expanding and connecting nodes in Cytoscape based on the graph structure in the database.

**Table 1:** Number of nodes in the database.

| Name of node | Total number |
| --- | --- |
| GO term | 47,059 |
| enzyme | 3,019 |
| pathway | 114 |
| map | 164 |
| compound | 4,328 |
|  | **54,684** |

**Table 2:** Number of relationships in the database.

| Name of relationship | Total number |
| --- | --- |
| crossConnect | 4,872 |
| isa | 71,400 |
| is | 228 |
| usedby | 11,248 |
| maplink | 5,300 |
| ECrel | 25,878 |
| in | 22,948 |
| produces | 12,150 |
| reaction | 8,032 |
|  | **162,056** |

### Mining the data using Cypher and Cytoscape

The final graph based on the biological data used (17 species) is a DCG containing in total 54,684 nodes (Table 1) and 162,056 relationships (Table 2). Using Cytoscape and the Neo4j plugin the graph can be queried and visualized to mine the functional annotation of the individual species and the four groups (Fig. 4). As an example, say that we would like to know which elements in the database are found in all species, which are specific for the culturable species, for all obligatory biotrophes, obligatory biotrophic chytrids and only higher fungi, or not found in the obligatory biotrophic chytrids. This information is extracted from the database with Cypher queries and by using the style filters in Cytoscape, each of these classes are assigned a different color (Fig. 5). Results show that the “Glyoxylate and dicarboxylate metabolism” pathway (path:ec00630) is missing 4 enzymes in the obligatory biotrophic chytrids (Query 2). Fig. 5 shows the compounds connected to, and the GO terms associated with, these 4 enzymes. Node colors indicate to which of the classes each element in the graph belongs.

**Figure 4:**
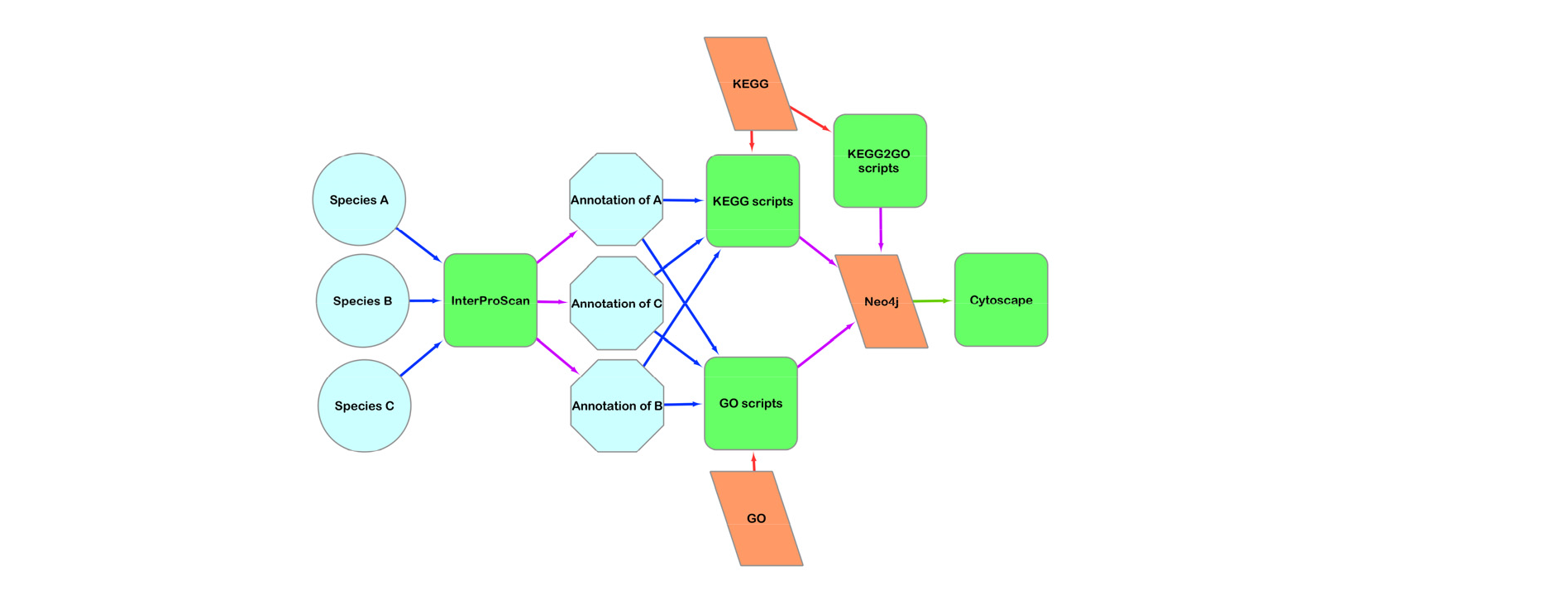
Flow of data for mining functional annotations. From left to right: the analyses start by annotating predicted proteins with InterProScan. Each of the annotation XML files is processed to link prestored KEGG and GO terms in the Neo4j database to the species by counting the number of occurrences. For KEGG annotations, the API is queried for additional information such as pathway names, compounds and GO terms linked to the enzymes. Hence only relevant information from KEGG is stored, not the entire database. Through Cytoscape’s Neo4j plugin specific subgraphs are queried, visualized and styled based on, for example, group information.

**Figure 5:**
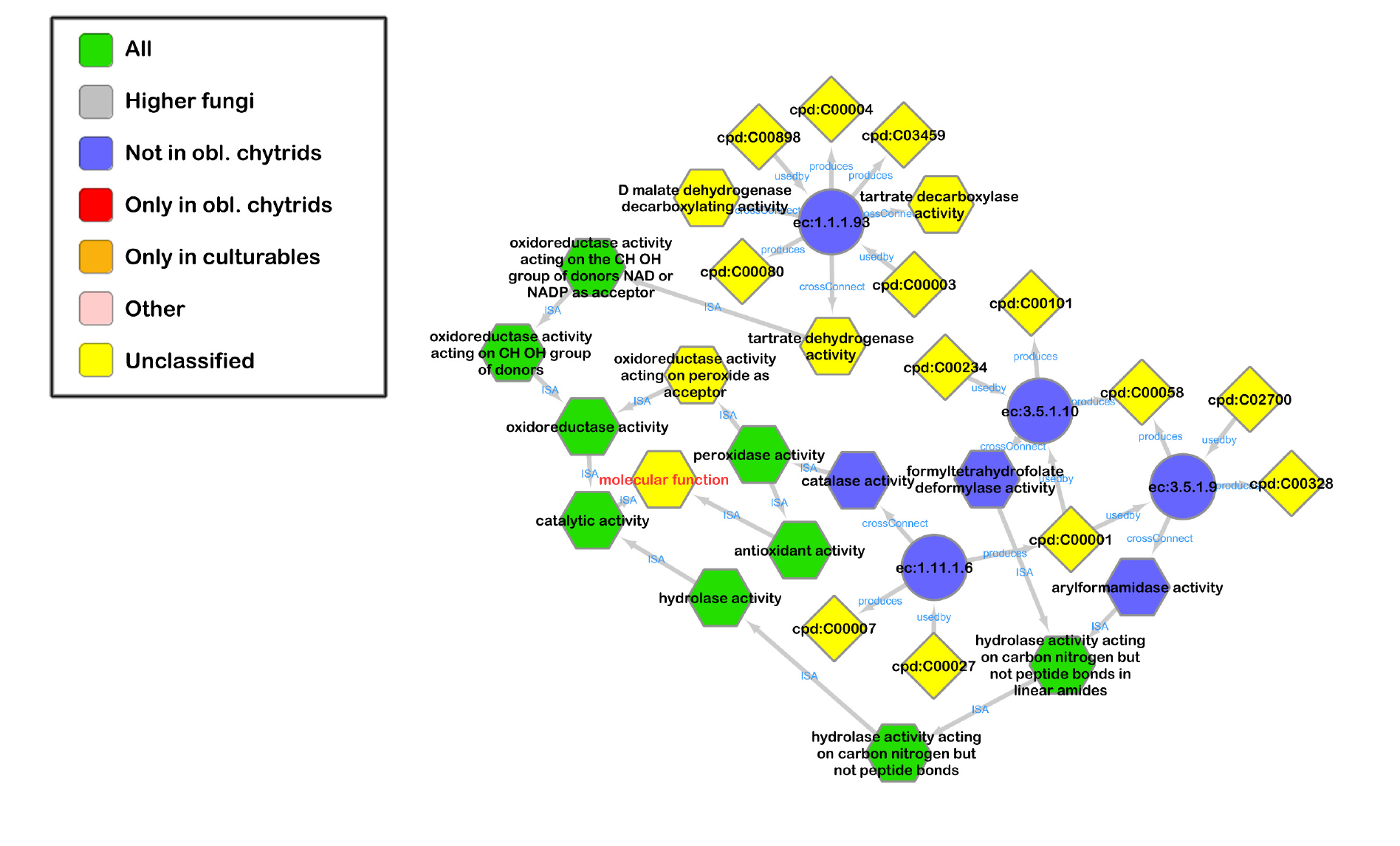
The four missing enzymes (circles) of pathway ec:00630 (“Glyoxylate and dicarboxylate metabolism”) in the two obligatory biotrophic chytrids and the compounds (diamonds) associated with the enzymes. GO terms linked to these enzymes are shown as hexagons with the paths to the toplevel node ‘molecular function’ (text in red).

~~~
   MATCH
   (p:pathway{name:'path:ec00630'})<-[:in]-(e:enzyme)
   WHERE e.group = 'noChytObl'
   WITH e
   OPTIONAL MATCH
   prod = (c1)-[:usedby]->(e)-[:produces]-(c2)
   OPTIONAL MATCH
   go=(g)<-[:crossConnect]-(e)
   WITH prod, go,g
   OPTIONAL MATCH
   fullGo=(headNode)<-[:ISA*]-(g)
   WHERE headNode.name = 'biological process' or headNode.name = 'cellular
   component' or headNode.name = 'molecular function'
   RETURN go, prod, fullGo
~~~

Query 2: Retrieve the missing enzymes in the obligatory biotrophic chytrids from the “Glyoxylate and dicarboxylate metabolism” pathway (path:ec00630), the compounds they are connected to, and the GO terms associated with these enzymes. Also trace back linked GO terms to the toplevel GO terms.

### Conclusion and discussion

The Neo4j graph database combined with the power of Cytoscape for visualization of selected parts of the data enables comparative analyses of GO, KEGG and InterProScan data from multiple genomes, either individually or grouped. For simplicity, the GO-basic dataset (with only “ISA” relationships) was used, making the graph less complicated than that of the full GO Plus dataset. This is not an intrinsic limitation: when the GO-plus dataset would be more appropriate, it can be used in the same way, although the resulting relationships are likely to become even more difficult to interpret. This manuscript focusses on the functional annotation of genes and does not take genome structure into account. To inspect genes associated with a particular enzyme or GO term a researcher has to visit an external genome browser. Some pangenome methods[23], such as PanTools[24], make it possible to store genome structures in a graph database. In the future, combining our approach with a pan genome graph will allow new ways of querying and visualizing multiple genomes. This will make it possible, for example, to retrieve the locations of genes used in a pathway of interest and extract sequence variability in these genes.

Another application of the presented approach is in metagenomics research. In environmental samples communities of organisms collaborate, consuming another organism’s products or compete for the same. The first step is to functionally annotate all genes found in the genome or transcriptome. The resulting graph can then be used to, for example, find out which pathways are present in the samples, which compounds are produced or which biological processes occur. This approach is not limited to a single metagenomics sample: by grouping annotations it is possible to compare samples to each other, as illustrated earlier by grouping multiple species.

The developed Cytoscape plugin is generic and not limited to the data sets presented here. Other publicly available databases such as Reactome[25] or any other Neo4j database can be queried through the plugin. An increasing number of bioinformatics tools, including for example tools for sequence alignment[26], support Neo4j, creating more demand for userfriendly graph visualization.

In this paper we showed that by combining and storing functional annotations of different species together with data from other sources, such as KEGG and GO, in Neo4j new approaches to species comparisons are now available. The Cytoscape plugin enable researchers to query these data and visualize them in userfriendly and powerful environment.

## Availability

The scripts, XML files and Neo4j database are available at https://github.com/swarris/sendo. The Neo4j plugin for Cytoscape is Open Source and available at the Cytoscape App Store (http://apps.cytoscape.org/apps/cytoscapeneo4jplugin) and at https://github.com/corwur/cytoscapeneo4j.

## Acknowledgements

We would like to thank Dick de Ridder and Jan-Peter Nap (both Wageningen UR) for their valuable comments on the manuscript. Ordina and Genetwister were so kind to allow SD and TvS work on the Open Source plugin on company time.

## Declarations

Ordina and Genetwister have no financial, legal or other benefits from this research and had no say in the content of the project. All authors declare no conflict of interest.

## Author contributions

SW wrote all Python scripts, performed the Interproscan data analysis, designed the cypher queries and created the plots. SW, SD and TvS redesigned, refactored and extended the Neo4j plugin for Cytoscape. BvdV supplied the biological data, created the groups of species and designed the biological experiments. SW and BvdV did the biological interpretation of the results. SW wrote the manuscript.

## References

1. Koren S, Walenz BP, Berlin K, Miller JR, Bergman NH, Phillippy AM. Canu: scalable and accurate long-read assembly via adaptive kmer weighting and repeat separation. Genome Res. 2017;27: 722–736. doi:10.1101/gr.215087.116

2. Qian Z, Adhya S. DNA repeat sequences: diversity and versatility of functions. Curr Genet. 2017;63: 411–416. doi:10.1007/s00294-016-0654-7

3. Hoff KJ, Lange S, Lomsadze A, Borodovsky M, Stanke M. BRAKER1: unsupervised RNA-Seq based genome annotation with GeneMark-ET and AUGUSTUS. Bioinformatics. 2015;32: 767– 769. doi:10.1093/bioinformatics/btv661

4. Holt C, Yandell M. MAKER2: an annotation pipeline and genome-database management tool for second-generation genome projects. BMC Bioinformatics. 2011;12: 491. doi:10.1186/1471210512491

5. Warris S, Boymans S, Muiser I, Noback M, Krijnen W, Nap J-P. Fast selection of miRNA candidates based on large-scale pre-computed MFE sets of randomized sequences. BMC Res Notes. 2014;7: 34. doi:10.1186/17560500734

6. Zdobnov EM, Apweiler R. InterProScan - an integration platform for the signaturere-cognition methods in InterPro. Bioinformatics. 2001;17: 847–848. doi:10.1093/bioinformatics/17.9.847

7. Gotz S, Garcia-Gomez JM, Terol J, Williams TD, Nagaraj SH, Nueda MJ, et al. High-throughput functional annotation and data mining with the Blast2GO suite. Nucleic Acids Res. 2008;36: 3420–3435. doi:10.1093/nar/gkn176

8. Kanehisa M, Goto S. KEGG: Kyoto Encyclopedia of Genes and Genomes. Nucleic Acids Res. 2000;28: 27–30. doi:10.1093/nar/28.1.27

9. de Vries RP, Riley R, Wiebenga A, Aguilar-Osorio G, Amillis S, Uchima CA, et al. Comparative genomics reveals high biological diversity and specific adaptations in the industrially and medically important fungal genus Aspergillus. Genome Biol. 2017;18: 28. doi:10.1186/s13059-017-1151-0

10. Carbon S, Ireland A, Mungall CJ, Shu S, Marshall B, Lewis S. AmiGO: online access to ontology and annotation data. Bioinformatics. 2009;25: 288–289. doi:10.1093/bioinformatics/btn615

11. Miller JJ. Graph database applications and concepts with Neo4j. Proceedings of the Southern Association for Information Systems Conference, Atlanta, GA, USA. 2013.

12. Shannon P, Markiel A, Ozier O, Baliga NS, Wang JT, Ramage D, et al. Cytoscape: a software environment for integrated models of biomolecular interaction networks. Genome Res. 2003;13: 2498–504. doi:10.1101/gr.1239303

13. Dijkxhoorn S, Sloten T van, Warris S. Cytoscape Neo4J Plugin [Internet]. 2017. Available: https://github.com/corwur/cytoscapeneo4j

14. Neo4J [Internet]. [cited 1 Sep 2016]. Available: https://neo4j.com/

15. Python. In: http://www.python.org [Internet]. Available: http://www.python.org

16. Cock PJA, Antao T, Chang JT, Chapman BA, Cox CJ, Dalke A, et al. Biopython: freely available Python tools for computational molecular biology and bioinformatics. Bioinformatics. 2009;25: 1422–3. doi:10.1093/bioinformatics/btp163

17. Kawashima S, Katayama T, Sato Y, Kanehisa M. KEGG API: a web service using SOAP/WSDL to access the KEGG system. Genome Informatics. 2003;14: 673–674. doi:10.11234/gi1990.14.673

18. Neo4J Team. Neo4J Python driver [Internet]. Available: https://github.com/neo4j/neo4jpythondriver

19. Browne F, Wang H, Zheng H. Investigating the impact human protein–protein interaction networks have on disease-gene analysis. Int J Mach Learn Cybern. 2018;9: 455–464. doi:10.1007/s1304201605035

20. Liu W, Li L, Long X, You W, Zhong Y, Wang M, et al. Construction and analysis of gene co-expression networks in Escherichia coli. Cells. 2018;7: 19. doi:10.3390/cells7030019

21. Summer G. cyNeo4j [Internet]. 2014. Available: https://github.com/gsummer/cyNeo4j

22. Warris S. SENDO Github page [Internet]. 2018. Available: https://github.com/swarris/sendo

23. Marschall T, Marz M, Abeel T, Dijkstra L, Dutilh BE, Ghaffaari A, et al. Computational pan-genomics: status, promises and challenges. Brief Bioinform. 2016;19: bbw089. doi:10.1093/bib/bbw089

24. Sheikhizadeh S, Schranz ME, Akdel M, de Ridder D, Smit S. PanTools: representation, storage and exploration of pan-genomic data. Bioinformatics. 2016;32: i487–i493. doi:10.1093/bioinformatics/btw455

25. Fabregat A, Korninger F, Viteri G, Sidiropoulos K, Marin-Garcia P, Ping P, et al. Reactome graph database: Efficient access to complex pathway data. PLOS Comput Biol. 2018;14: e1005968. doi:10.1371/journal.pcbi.1005968

26. Warris S, Timal NRN, Kempenaar M, Poortinga AM, van de Geest H, Varbanescu AL, et al. pyPaSWAS: Python-based multicore CPU and GPU sequence alignment. PLoS One. 2018;13: e0190279. doi:10.1371/journal.pone.0190279

